# Video-rate volumetric functional imaging of the brain at synaptic resolution

**DOI:** 10.1101/058495

**Authors:** Rongwen Lu, Wenzhi Sun, Yajie Liang, Aaron Kerlin, Jens Bierfeld, Johannes Seelig, Daniel E. Wilson, Benjamin Scholl, Boaz Mohar, Masashi Tanimoto, Minoru Koyama, David Fitzpatrick, Michael B. Orger, Na Ji

## Abstract

Neurons and neural networks often extend hundreds to thousands of micrometers in three dimensions. To capture all the calcium transients associated with their activity, we need volume imaging methods with sub-second temporal resolution. Such speed is challenging for conventional two-photon laser scanning microscopy (2PLSM) to achieve, because of its dependence on serial focal scanning in 3D and the limited brightness of indicators. Here we present an optical module that can be easily integrated into standard 2PLSMs to generate an axially elongated Bessel focus. Scanning the Bessel focus in 2D turned frame rate into volume rate and enabled video-rate volumetric imaging. Using Bessel foci designed to maintain synaptic-level lateral resolution *in vivo*, we demonstrated the power of this approach in enabling discoveries for neurobiology by imaging the calcium dynamics of volumes of neurons and synapses in fruit flies, zebrafish larvae, mice, and ferrets *in vivo*.

## INTRODUCTION

Understanding neural computation requires the ability to measure how a neuron integrates its multitude of synaptic inputs, as well as how neural ensembles work together during sensory stimulation and motor control. With submicron lateral resolution and optical sectioning ability in scattering brains, two-photon laser scanning microscopy^1, 2^ (2PLSM) has become a powerful tool for monitoring the activity of neurons and their networks *in vivo* through their calcium transients. However, both individual neurons and neural networks may extend over hundreds or even thousands of micrometers in three dimensions. Because conventional 2PLSM images volume by scanning the excitation laser focus serially in 3D, the limited brightness of calcium indicators and the inertia of laser scanning units make it difficult to capture all the calcium transients in a volume at video rate (e.g., 30 Hz) while maintaining synapse-resolving ability.

Realizing that for most *in vivo* brain imaging experiments, the positions of neurons as well as their synapses are known and remain unchanged during each imaging session, one can increase volume imaging speed substantially by sacrificing axial resolution. By scanning an axially elongated focus (e.g., a Bessel beam) in 2D^3–6^, we obtained projected views of 3D volumes and converted 2D frame rates into 3D volume rates (**Fig. 1a**). Utilizing a spatial light modulator, we built a module that generated Bessel foci optimized for *in vivo* brain imaging. Easily incorporated into three different 2PLSM systems, this simple module allowed volume imaging rates of up to 30 Hz while maintaining the ability to resolve dendritic spines and axonal boutons. To demonstrate its versatility, we applied the Bessel focus scanning method to imaging brain volumes of a variety of model systems *in vivo*, such as fruit fly, zebrafish larva, mouse, and ferret. High speed volume imaging was used to inform on the tuning properties of dendritic spines and the synchrony of inhibitory neuron activity in mouse visual cortex, the network dynamics of reticulospinal neurons in the hindbrains of zebrafish larvae, and the responses of fruit fly brains to visual stimuli.

## RESULTS

### Optimized Bessel foci enable 30 Hz *in vivo* volume imaging of dendrites, axons, and synapses

The volume-imaging module was composed of a phase-only spatial light modulator (SLM) and an annular mask array, and was incorporated into a 2PLSM by placing it between the excitation laser and the microscope (**Fig. 1b**). A concentric phase pattern on the SLM generated a ring pattern at the focal plane of lens L1. After spatial filtering with an annular mask, the ring pattern was imaged onto the galvanometer scanners by lenses L2 and L3. This led to an annular illumination on the back pupil plane of the microscope objective and an axially extended focus approximating a Bessel beam. Combining different SLM patterns with distinct annular masks, we generated a series of Bessel foci with various axial full widths at half maximum (axial FWHM, 15 to 400 μm) and numeric apertures (NA, 0.4 to 0.93) (**Methods**), and characterized their performance for *in vivo* brain imaging on three 2PLSM systems (one commercial and two homebuilt systems; see **Supplementary Table 1** for focal lengths of L1-3 for these three systems). We discovered that lower NA Bessel foci afford better *in vivo* image quality than those with higher NA (**Supplementary Fig. 1**). This is because high-NA Bessel foci have too much energy distributed in their side rings, which, even with suppression by nonlinear excitation, still leads to substantial background signal and causes image blur. Because the central peak of a Bessel focus is sharper than that of a Gaussian focus^3, 7^, even at NA 0.4, a Bessel focus still has sufficient lateral resolution to resolve individual synaptic terminals but generates much less background haze. Therefore, for all subsequent experiments, we employed Bessel foci with NA 0.4.

**Figure 1.**
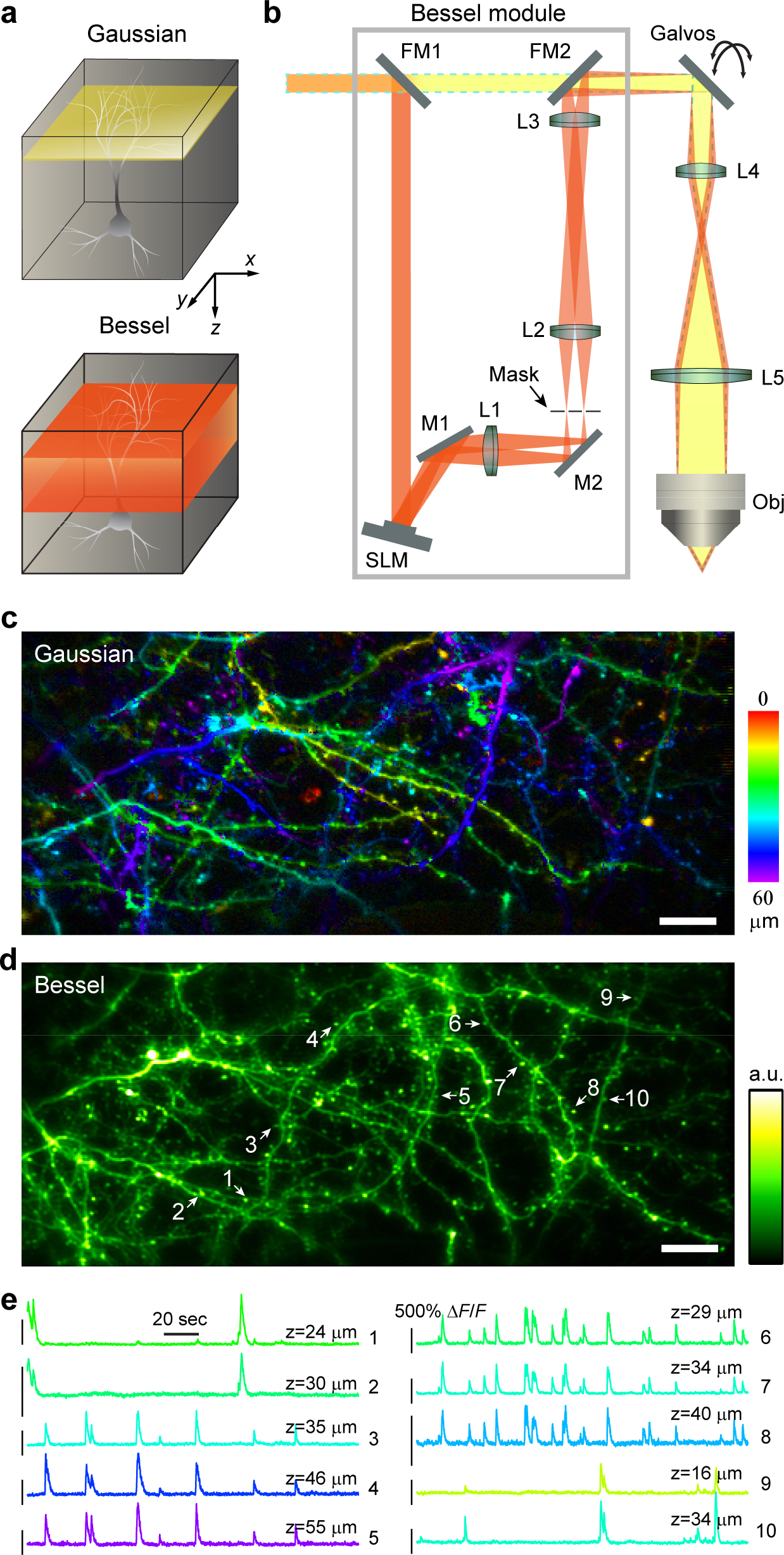
Concept, design, and performance of Bessel module for *in vivo* volume imaging. (**a**) Scanning a Gaussian focus in *xy* plane images a thin optical section (yellow shaded region), whereas scanning an elongated Bessel focus in *xy* informs on the structures in a 3D volume (red shaded region). (**b**) Schematics of Bessel module, which is composed of a spatial light modulator (SLM), three lenses (L1, L2, L3), and an annular mask. The module generates an annular illumination pattern on the galvanometers (Galvo) and, after conjugation by lenses L4 and L5, at the back pupil of the objective (Obj), resulting in a focus approximating a Bessel beam. M1 and M2: folding mirrors; FM1 and FM2: flip mirrors to switch between Gaussian (yellow with dashed blue outline) and Bessel (red) paths. (**c-e**) Example application of Bessel scanning to imaging a volume of GCaMP6s^+^ neurites in awake mouse cortex at 30 Hz. (**c**) Mean intensity projection of a 60-μm-thick image stack collected with conventional Gaussian focus scanning, with structures color-coded by depth. (**d**) Image of the same volume of neurites collected by scanning a Bessel focus with 0.4 NA and 53 μm axial FWHM. Arrows and numbers labels individual axonal varicosities (putative boutons, 1,2) and dendritic spines (3-10). (**e**) Representative calcium transients measured in boutons and spines from four neurons (color-coded by *z* depth). Objective: Olympus 25×, 1.05 NA; Wavelength: 960 nm; Post-objective power: 30 mW for Gaussian and 118 mW for Bessel scanning. Scale bar: 20 μm. Representative images from 3 mice are shown.

**Supplementary Figure 1.**
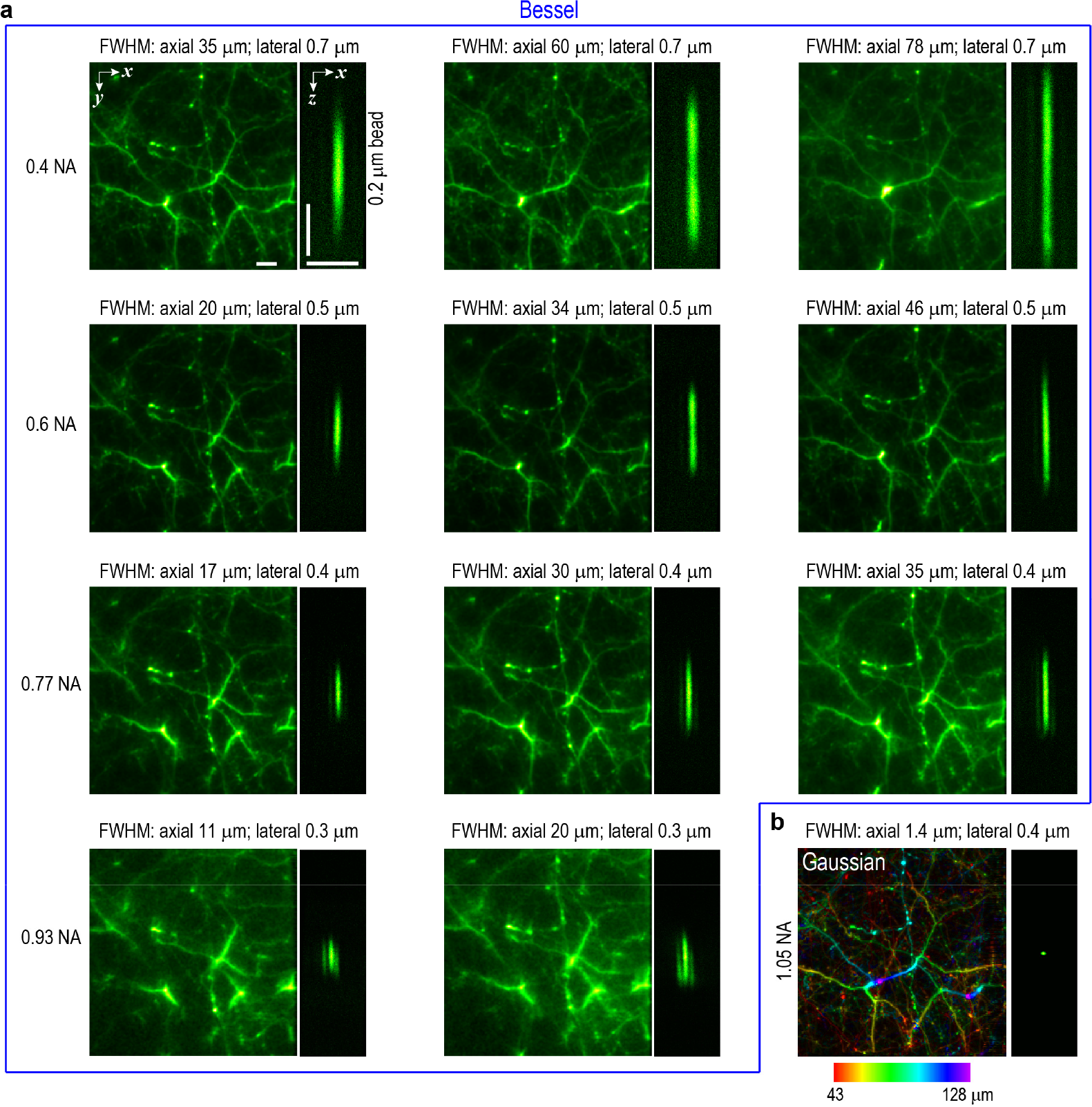
Optimize Bessel foci for *in vivo* volume imaging. (**a**) Images taken by scanning Bessel foci of various NAs, lateral and axial FWHMs: (Left panels) *in vivo* volume images of YFP^+^ neurites in cortex of an awake mouse (*Thy1*-YFP); scale bar: 10 μm. (Right panels) Axial point spread functions measured from 0.2-μm-diameter beads; ***x* scale bar: 5 μm; *z* scale bar: 25 μm.** (**b**) Mean intensity projection of the same neurites (color-coded by depth below pia) imaged by scanning a 1.05-NA Gaussian focus in 3D. Note the similarity in quality between the 0.4/0.6-NA Bessel foci and **b**. Higher NA Bessel foci have stronger side rings, resulting in hazy backgrounds. Objective: Olympus 25×, 1.05 NA.

To demonstrate the speed and resolution of Bessel volume scanning, we imaged sparse GCaMP6s^+^ neurites (0-60 μm below pia) in the primary visual cortex (V1) of awake mice. With a Gaussian focus (NA 1.05), conventional 3D scanning (60 2D frames spaced by 1 pm in axial direction) was required to image a volume extending 60 μm in *z* (**Fig. 1c**). In contrast, scanning a Bessel focus (NA 0. 4, axial FWHM 53 μm, **Fig. 1d**) in 2D captured all the structures in one frame, reducing the data size ~60 fold. Using a 2PLSM with a resonant galvo, visually-evoked calcium transients in axonal varicosities
and dendritic spines within a 270μm×270μm×60μm volume were measured at 30 Hz (**Supplementary Video 1, Fig. 1e**). Importantly, with Bessel scanning, the gain in volume imaging speed does not come with the reduction of lateral resolution, with synapses (arrows in **Fig. 1d**) clearly resolvable even in single frames taken at 30 Hz (**Supplementary Fig. 2**).

One unique strength of Bessel scanning as a volumetric imaging tool is its robustness against artifacts induced by brain motion. In awake behaving mice, the brain may move both laterally (within the *xy* plane) and axially (along *z*). Whereas lateral motions can be corrected post hoc^8, 9^, axial motion correction requires one to constantly image volumes encompassing the entire *z* motion range^10^, which may vary from 2 μm (in headfixed running mice^8^) to 15 μm (induced by licking^10^). Because axial lengths of Bessel foci can be substantially larger than typical ranges of axial brain movement, motion along *z* affects Bessel-scan images minimally. In other words, because a Bessel focus is basically light whose lateral spread (as determined by its NA or lateral resolution) remains small over a large axial propagation distance (as indicated by the axial FWHM), during Bessel focus scanning, a volume is imaged constantly with the axial scan speed being the speed of light. As a result, no postprocessing is needed for registration in *z*. As a demonstration, using a extended Bessel focus (NA 0.4, axial FWHM 91 μm), we imaged at 30 Hz dendritic branches (180μm× 180μm × 160μm) of a GCaMP6f^+^ neuron in the somatosensory cortex (S1) of an awake mouse. Despite substantial lateral movement, the entire structure stayed within the extended Bessel focus (**Supplementary Video 2**). Simple 2D image registration using TurboReg plugin of ImageJ^11, 12^ was sufficient to register these 30 Hz time-series of 180μm× 180μm × 160μm volume (**Supplementary Fig. 3, Supplementary Video 2**).

**Supplementary Figure 2.**
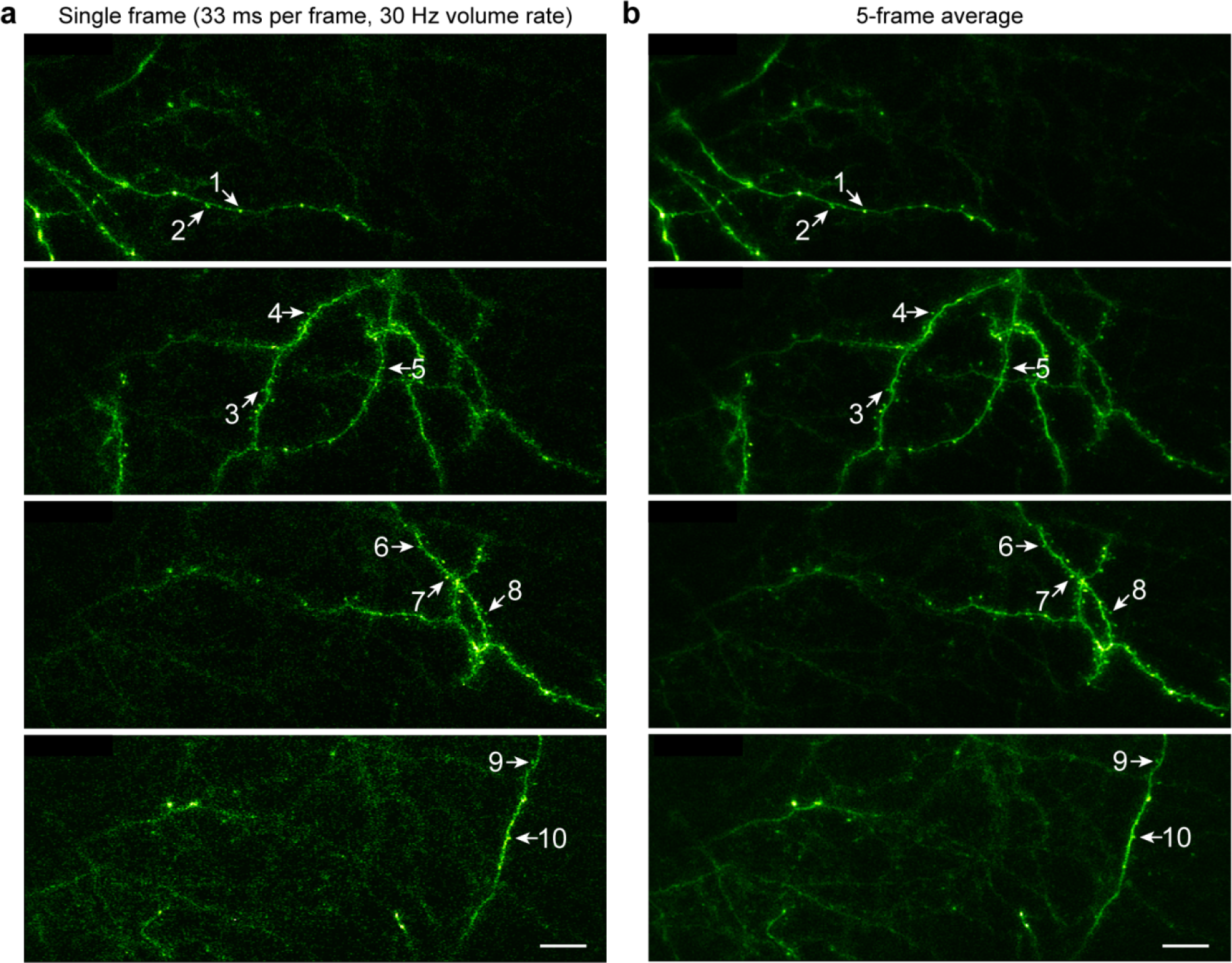
Axonal varicosities and dendritic spines are resolvable in single frames of 30 Hz Bessel volume scanning. (**a**) Individual raw image frames during 30-Hz volumetric measurements of calcium transients in a volume extending 60 μn in *z*. Arrows and numbers label individual axonal varicosities (putative boutons, 1,2) and dendritic spines (3-10) (same as in **Fig. 1d**). (**b**) As comparison, the same synaptic structures in the average of 5 frames (i.e., effective volume rate of 6 Hz). Scale bars: 20 μm.

**Supplementary Figure 3.**
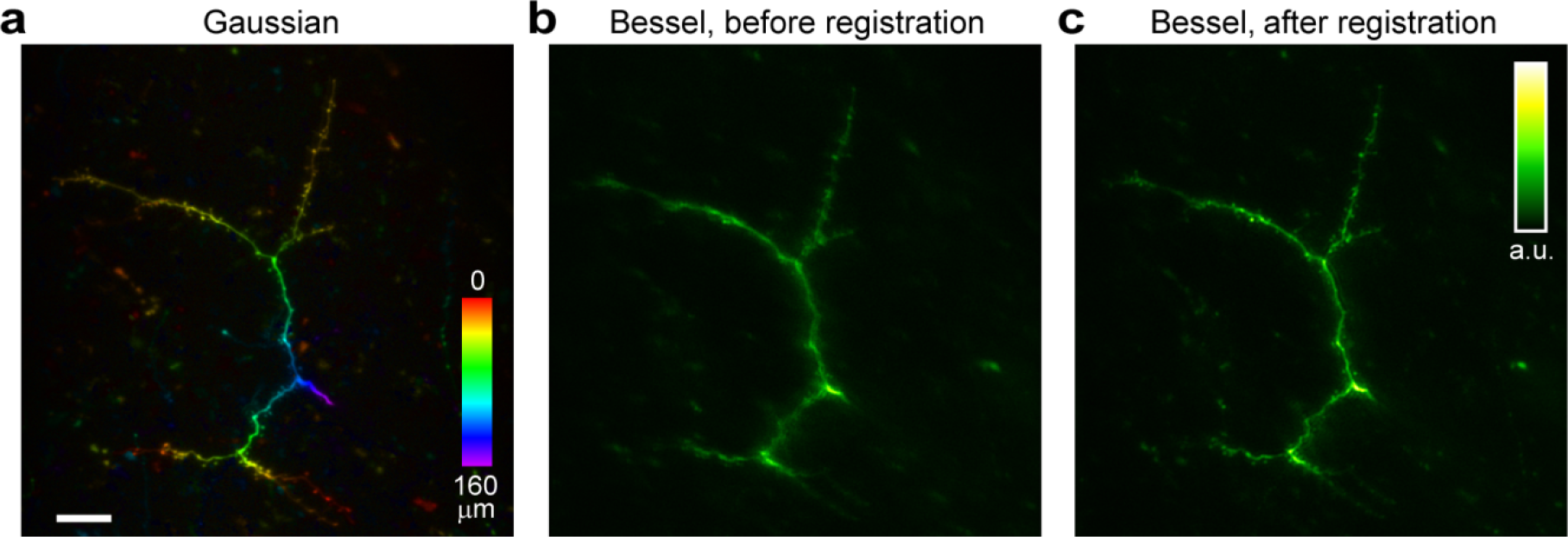
Bessel scanning data only require 2D registration. (**a**) Mean axial projection of a 3D image stack (180μm× 180μm × 160μm, 100 2D images) collected by Gaussian focus scanning, showing dendritic branches (0-160 μm below pia) in S1 of an awake mouse, with structures color-coded by depth. (**b**) Averaged image of a 266-sec time series of the same dendrites imaged with a Bessel focus (0.4 NA and 91 μm axial FWHM) at 30 Hz. (c) Averaged image after registering the time series using TurboReg plugin of ImageJ. Objective: Olympus 25×, 1.05 NA; Post-objective power: 30 mW for Gaussian focus scanning and 103 mW for Bessel focus scanning; Wavelength: 960 nm; Scale bars: 20 μm. (Also see **Supplementary Video 2**).

### High-throughput volumetric functional mapping of single spines in mouse and ferret V1

Having demonstrated the feasibility of rapid volumetric imaging of dendritic spines using Bessel foci, we applied this technique to high-throughput functional mapping of sensory-evoked calcium transients in single spines. Because of their small sizes and three dimensionally distributed locations on dendrites, imaging dendritic spines over large volumes *in vivo* is technically challenging^13^. With conventional Gaussian focus scanning, multiple imaging planes need to be acquired and tiled to cover a single branch^14, 15^. With the extended depth of focus offered by Bessel foci, we could image calcium transients in dendritic spines along multiple dendritic branches simultaneously.

We first investigated the tuning properties of spines on a L2/3 neuron in V1 of an anesthetized mouse presented with drifting grating stimuli (12 drifting directions presented with pseudorandom sequences for 10 trials). Conventional Gaussian focus scanning in 3D provided a depth-resolved image stack of GCaMP6s^+^ spines along dendritic branches 100-160 μm below pia (**Fig. 2a**). Scanning a Bessel focus (NA 0.4, axial FWHM 35 μm) in 2D, we resolved these spines in a 102μm×102μm× 60μm volume (**Fig. 2b**) and measured their visually evoked spine-specific calcium transients at 1.5 Hz (**Supplementary Video 3**). Following previous work^15^, we removed transients induced by the occasional back-propagating action potential (BAP) from the calcium signal, and quantified the spine calcium transients as percentage fluorescence change Δ*F/F* (Δ*F*=*F*_response_−*F*_baseline_, *F*=*F*_baseline_). Of all 75 identified dendritic spines, 57.3% (43 spines) exhibited visually evoked activity (Δ*F/F*>10%), with 36 spines also showing significant tuning to grating angles (*P*<0.01, ANOVA across 12 directions, see **Methods**). Representative spines (s1-11 in **Fig. 2**) showed large visually evoked transients selective to grating angle (**Figs. 2c,d**), but independent of their neighboring dendritic shafts (d1-11 in **Fig. 2**), which showed little responses with no significant tuning preference (**Fig 2c**). We found that spines on the same branch could have different tuning preferences (**Figs 2e,f**), and the tuning of the BAP-induced calcium transients in dendritic shafts (indicative of somatic tuning) was similar to the histogram of the preferred grating angles of all tuned spines (**Fig. 2e**), consistent with previous observations^14, 15^. We succeeded, within one day, in integrating a Bessel scanning module into a 2PLSM system that was designed to image ferret brains^16^. We then measured calcium transients from spines distributed over 72 μm axial range within ferret V1 at 30 Hz and determined their tuning towards drifting gratings (**Supplementary Fig. 4**).

**Figure 2.**
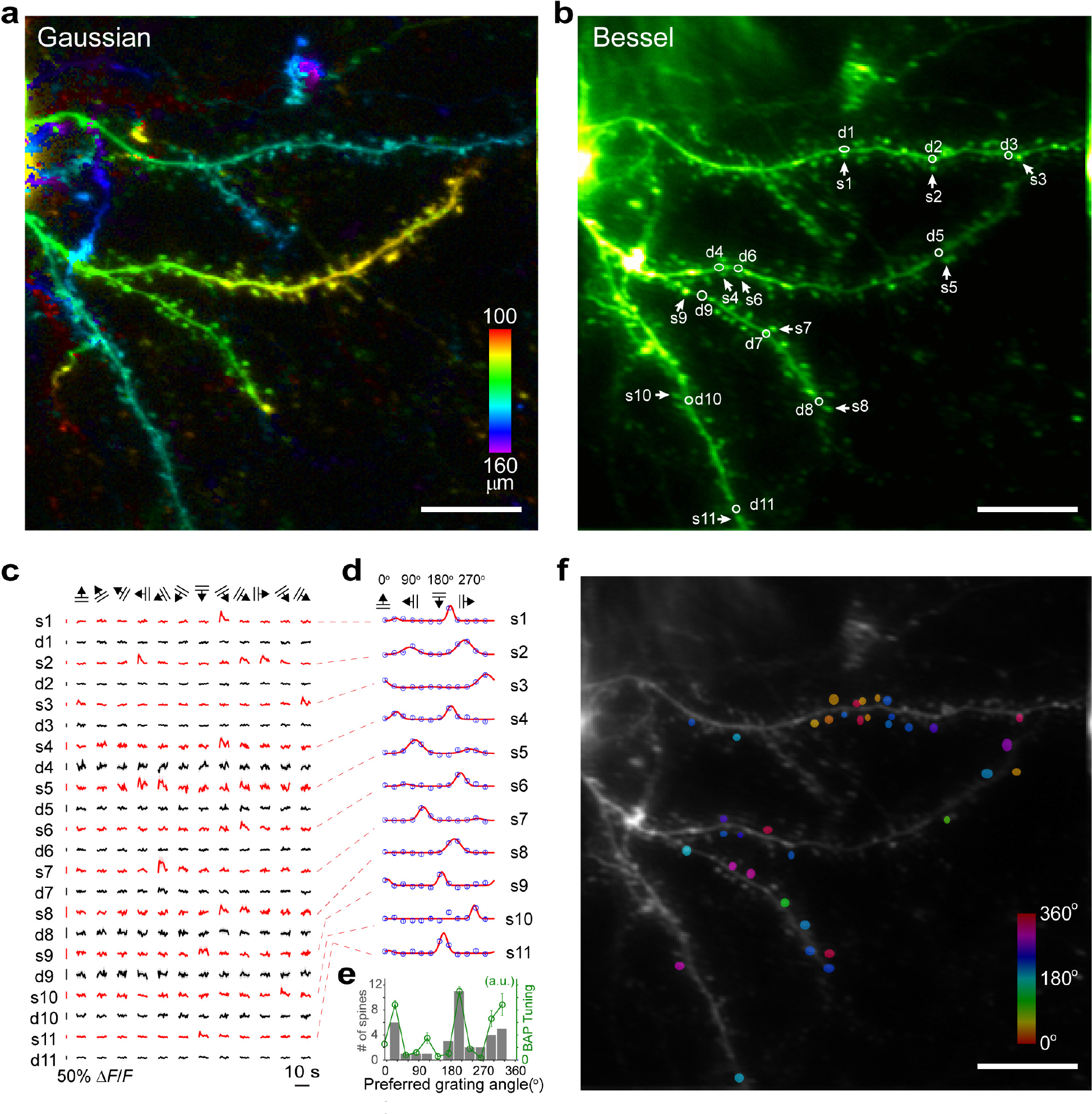
High-throughput *in vivo* characterization of orientation tuning of dendritic spines in mouse V1. (**a**) Mean intensity projection of a 3D image stack (102μm×102μm×60μm) collected by Gaussian focus scanning of dendritic branches (100-160 μm below pia) in V1 of an anesthetized mouse, with structures color-coded by depth. (**b**) Image of the same branches collected by scanning a Bessel focus with 0.4 NA and 35 μm axial FWHM. Example spines (arrows, s1-s11) and neighboring dendritic shafts (circles, d1-d11) are indicated. (**c**) 10-trial-averaged calcium transients of spines (red) and neighboring dendritic shafts (black) in b for 12 grating angles. (**d**) Tuning curves of individual spines calculated from c. (**e**) (Gray histogram) distribution of preferred grating angles of all tuned spines and (green line and symbol) tuning curve of BAP-induced calcium transient in dendritic branches. (All four main branches had similar tuning curves. Only one was shown.) (**f**) Locations of all tuned spines highlighted with color overlay, with the colors indicating preferred grating angles. Objective: Olympus 25×, 1.05 NA; Post-objective power: 33 mW for Gaussian scanning and 74 mW for Bessel scanning; Wavelength: 960 nm; Scale bars: 20 μn; Shadow (**c**) and error bars (**d,e**) represent s.e.m. Representative results from 3 mice.

**Supplementary Figure 4.**
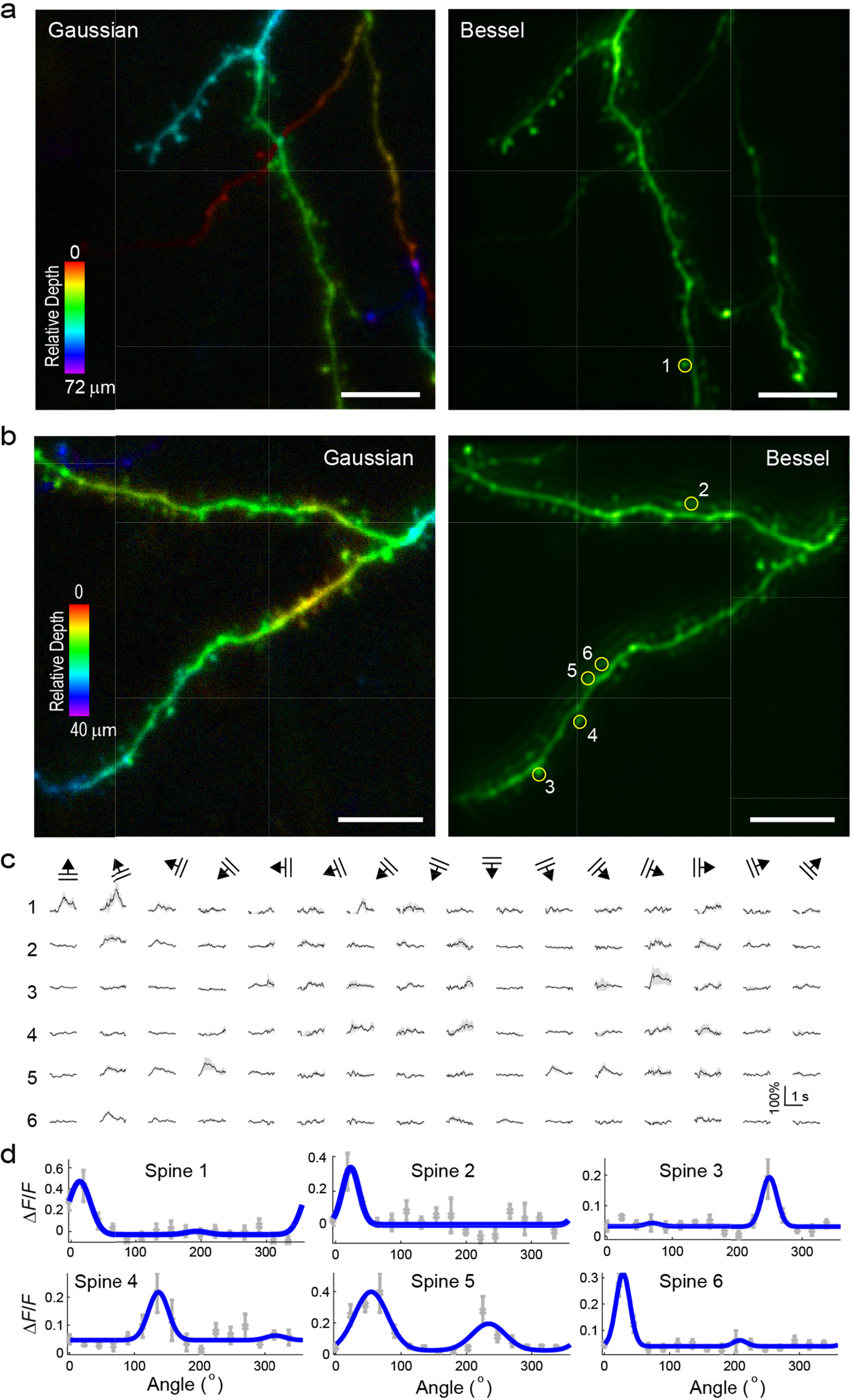
*In vivo* characterization of dendritic spines in ferret V1 at 30 Hz volume rate. (**a,b**) Two examples of applying Bessel focus scanning to the characterization of spines in ferret V1. (Left) Mean intensity projection of 3D image stacks by conventional Gaussian focus imaging in V1 of anesthetized ferrets; (Right) Images collected by scanning a Bessel beam with 0.4 NA and 50 μm axial FWHM. Yellow circles highlight example spines. (**c**) 10-trial-averaged calcium transients of spines in **a** and **b** for 12 grating angles. (**d**) Tuning curves of individual spines in **a** and **b**. Objective: Nikon 16×, NA 0.8; Post objective power: 55 mW for Gaussian scanning and 84 mW for Bessel scanning; Wavelength: 960 nm; Shadow (**c**) and error bars (**d**): s.e.m; Scale bars: 10 μm. Representative images from 2 ferrets.

### Population dynamics of inhibitory neuron ensembles in V1 of awake mice

In addition to monitoring the activity of neuronal compartments, Bessel focus scanning can also be used to monitor the population dynamics of 3D neural ensembles such as cortical GABAergic neurons. Representing 15-20% of all neurons in neocortex and playing important roles in shaping the functional architecture of cortical networks^17, 18^, these typically inhibitory neurons have been studied with 2PLSM *in vivo*^19–21^. Improved labeling strategies (through transgenic mouse strains^22–27^ or post hoc identification via immunohistochemistry^28^) have enabled imaging of specific subtypes of GABAergic neurons *in vivo*. However, the ensemble dynamics of GABAergic neurons are much less studied. One technical challenge for studying cell-type-specific ensemble activity is caused by their low density in neocortex: a typical 2D image plane may contain <10 GABAergic neurons ^19, 21, 28, 29^ and even fewer neurons of specific subtypes^22, 24, 25, 28^, making it difficult to measure the activity of large numbers of neurons at the same time. Using an extended Bessel focus (NA 0.4, axial FWHM 78 μm), we simultaneously captured the calcium transients of up to 71 GCaMP6s^+^ GABAergic neurons scattered within 256 μm × 256 μm × 100 μm volumes (0-100, 101-200, 201-300, 301-400 μm below pia) in V1 of quietly awake Gad2-ires-Cre mice^23^. Due to their low density, almost all neurons visible in the Gaussian 3D stacks could be identified in the projected volume view offered by Bessel focus scanning (**Figs. 3a,b**).

We measured how these GABAergic neurons responded to drifting grating stimuli. During experiments, we also monitored the pupil of the mouse, whose size fluctuation reflected brain state^26, 30, 31^. After subtracting the neuropil background^32^ from GCaMP6s^+^ neurites (**Methods, Supplementary Fig. 5**), we found, at all depths, that a substantial fraction of neurons were not visually driven, but had calcium transients strongly correlated with the pupil size and with each other (**Figs. 3c,d**). Such activity synchrony persisted in darkness (**Supplementary Fig. 6**) and was more widespread for superficial GABAergic neurons than for deeper ones. Calculating Pearson’s correlation coefficient *R* for pairs of neurons, we found that GABAergic neurons 0-200 μm below pia (*n*=348) had 41% of their *R*’s larger than 0.4, whereas neurons 201-400 μm below pia (*n*=383) only had 22% of all *R*’s larger than 0.4 (**Fig. 3e**).

**Figure 3.**
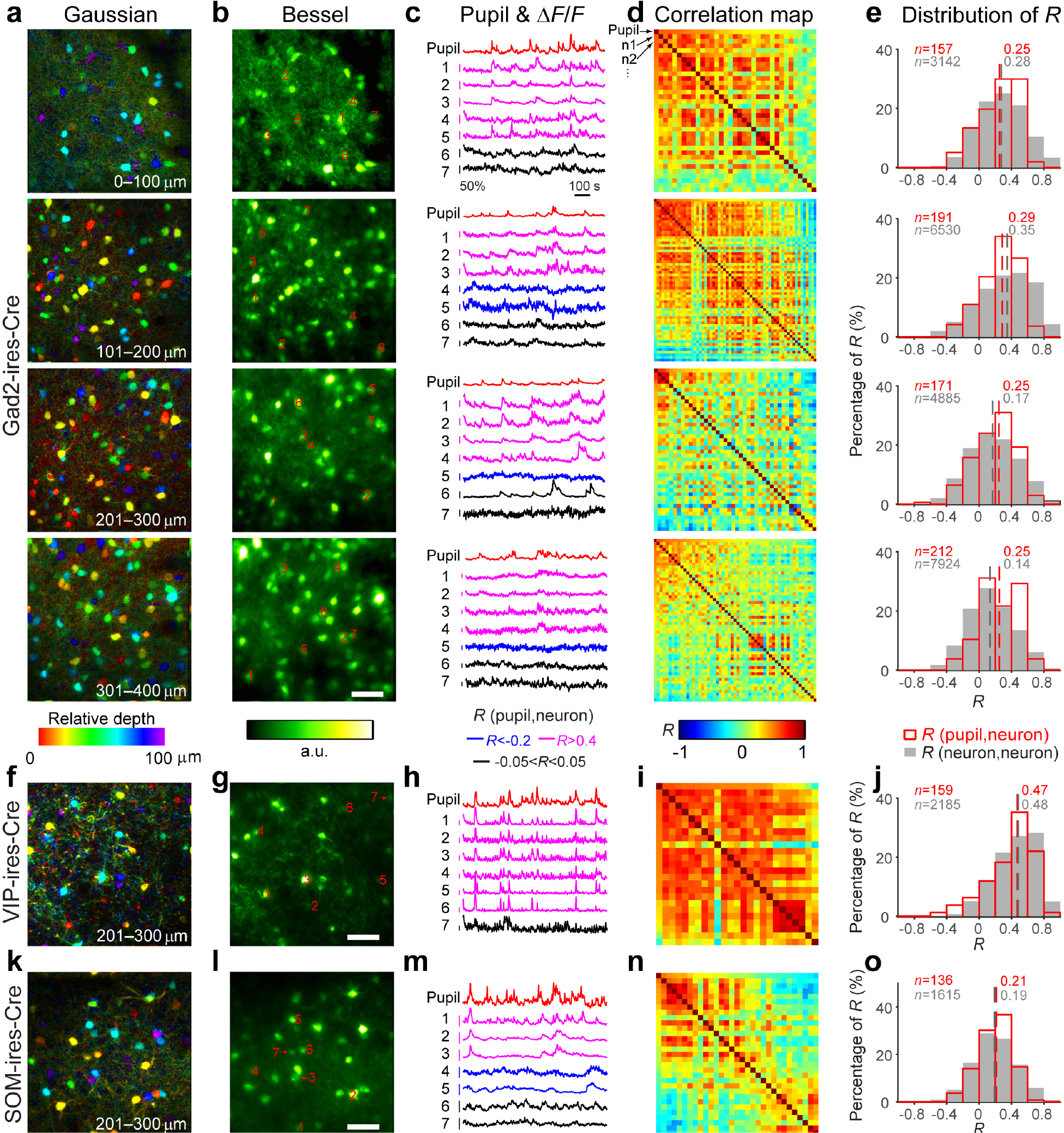
Activity synchrony among GABAergic neural ensembles in V1 of awake mice. (**a-e**) Measurements on Gad2-ires-Cre mice transfected with AAV-Syn-Flex-GCaMP6s. (**a**) Mean intensity projections of 3D volume stacks (256μm×256μm×100μm each, 0-100, 101-200, 201-300, 301-400 μm below pia, total 400 2D images) collected by scanning a Gaussian focus in V1, with neurons color-coded by depth. Post-objective power: 5, 5, 11, and 14 mW, respectively. (**b**) Images of the same neurons collected by scanning a Bessel focus with 0.4 NA and 78 μm axial **FWHM**. Post-objective power: 21, 25.2, 31.5, and 42 mW, respectively. (**c**) Pupil size (red) and concurrently measured calcium transients of neurons highlighted in **b** (blue if Pearson’s correlation coefficient *R*<−0.2, black if −0.05<*R*<0.05, magenta if *R*>0.4). (**d**) *R* between pupil size and calcium transients (1^st^ row and 1^st^ column, neurons sorted in descending order of *R*), and between the calcium transients of all pairs of neurons identifiable in **b**. (**e**) Distribution of *R* within four depth ranges measured from 3 mice. *n*: number of *R*’s; Red open histogram: *R* between pupil size and neuronal calcium transients; Gray filled histogram: *R* between neurons; Dashed lines: median. (**f-o**) Measurements and analysis on (**f-j**) VIP-ires-Cre and (**k-o**) SOM-ires-Cre mouse transfected with AAV-Syn-Flex-GCaMP6s. Histograms in **j** and **o** include all VIP^+^ and SOM^+^ neurons 1-400 μm below pia. Objective: Olympus 25×, 1.05 NA; Wavelength: 960 nm; Scale bar: 50 μm.

**Supplementary Figure 5.**
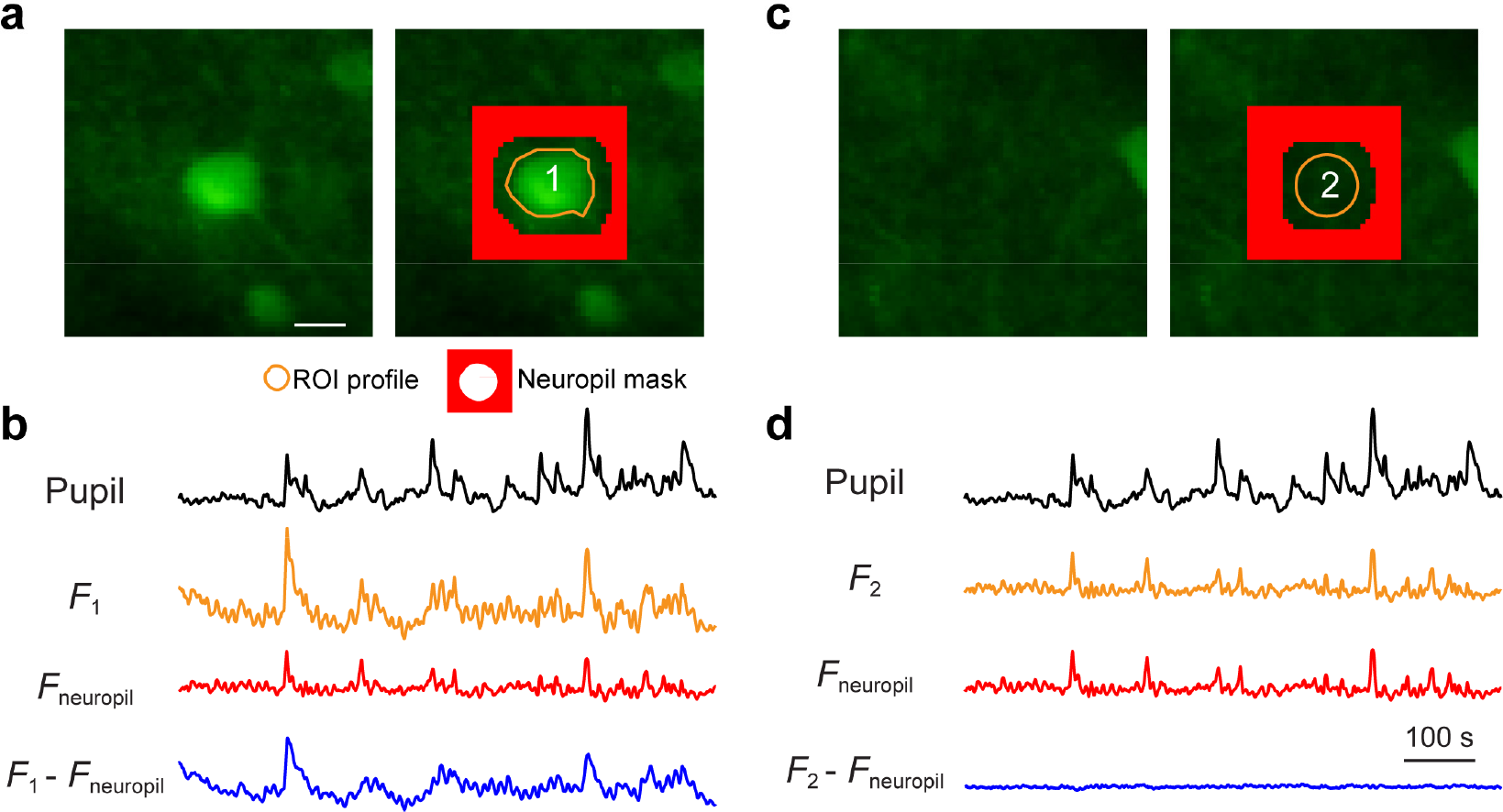
Removal of neuropil contamination in Bessel focus scanning mode. (**a**) (left) Image of a cell body obtained by scanning a Bessel beam with 0.4 NA and 78 μm axial FWHM, (right) overlaid with region of interest (ROI, orange outline) and the area used for calculating neuropil background (red region). (**b**) The difference between the averaged signal within ROI (*F*_1_) and the averaged signal within neuropil mask (*F*_neuropil_) is used to calculate calcium transient of the neuron. After the removal of neuropil contamination, the cell body still exhibited activity correlated with pupil size. (**c**) and (**d**) same as **a** and **b**, except that the removal of neuropil contamination was applied to a volume of neuropil, resulting in minimal calcium transients. All fluorescence traces were plotted on the same scale.

**Supplementary Figure 6.**
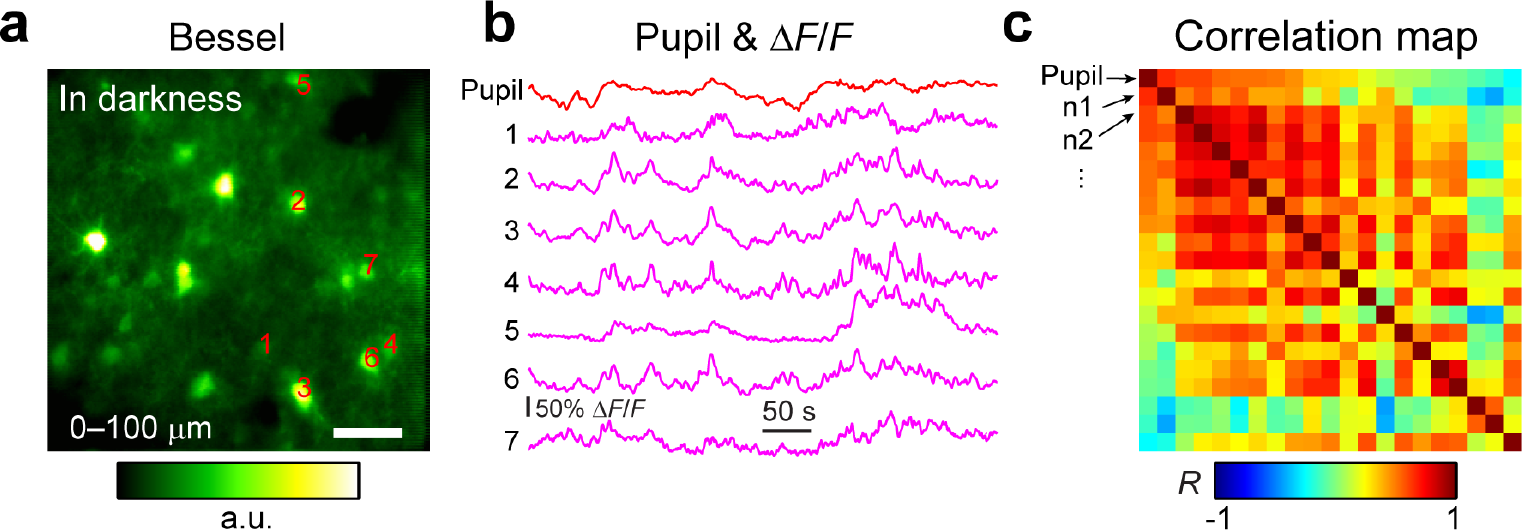
Activity synchrony among GABAergic neurons persists in the dark. (**a**) Representative image of GAD2^+^ neurons expressing GCaMP6s (0-100 μm below pia) collected by scanning a Bessel beam with 0.4 NA and 78 μm axial **FWHM**. (**b**) Pupil size (red) and concurrently measured calcium transients of example neurons highlighted in a. (c) Pearson’s correlation coefficient (*R*) between neuronal calcium transients and pupil size (1^st^ row and 1^st^ column, sorted by decreasing *R*), as well as between the calcium transients of all pairs of neurons identified in **a**. Objective: Olympus 25×, 1.05 NA; Post-objective power: 21 mW; Wavelength: 960 nm; Scale bar: 50 μm.

Because GABAergic neurons in cortex include several subtypes with distinct properties^17, 18^, we investigated how activity synchrony depended on cell type. We focused on two subtypes that were immuno-reactive for vasoactive intestinal peptide (VIP) and somatostatin (SOM), respectively, because they were known to be involved in cortical state changes as indicated by pupil size^26, 33^. Using transgenic mice^23^ (VIP-ires-Cre and SST-ires-Cre) where only VIP^+^ or SOM^+^ neurons expressed GCaMP6s, we imaged ensembles of VIP+ and SOM+ neurons using Bessel scanning during visual stimulus presentation (**Figs. 3g,l**, see **Figs. 3f,k** for Gaussian 3D stacks). We found that VIP^+^ neurons (*n*=159) had highly synchronized activity that was positively correlated with pupil size (60% of *R*’s larger than 0.4, **Figs. 3h-j**). In contrast, responses of SOM^+^ neurons (*n*=136) were more heterogeneous, and on the population level, less synchronized with pupil size (22% of *R*’s larger than 0.4, **Figs. 3m-o**).

These observations are in general agreement with known properties of VIP^+^ and SOM^+^ neurons. Earlier *in vivo* studies^25, 26^ found that individual VIP^+^ neurons became active during heightened attention/pupil dilation, consistent with our observation of the large *R*’s between neuronal calcium transients of VIP^+^ neurons and pupil size (**Figs. 3i,j**). Bessel scanning enabled us to obtain, to our knowledge, the first experimental evidence that VIP^+^ neuron ensembles in cortical volumes have highly synchronized, brain-state-correlated activity (possibly mediated by gap junctions^34, 35^). Given several recent studies showing that VIP^+^ neurons directly inhibit SOM^+^ neurons, we expected to see mostly negative correlation between SOM^+^ neurons and pupil size. However, we found SOM^+^ neurons to exhibit heterogeneous responses, with some negatively correlated with pupil size (and thus VIP^+^ neuron activity) while others positively correlated with pupil size (**Figs. 3m-o**). This is consistent with recent *in vivo* studies^26, 36^, where SOM^+^ neurons were found to have heterogeneous population responses including those co-active with VIP^+^ neurons. It is also consistent with the suggestion that SOM^+^ neurons in SST-ires-Cre mice may include multiple subtypes of inhibitory neurons^37^. The distinct activity patterns of VIP^+^ and SOM^+^ neurons may explain the decrease in activity synchrony of GABAergic neurons with depth (**Figs. 3d,e**). Because VIP^+^ neurons are more abundant in superficial layers whereas SOM^+^ neurons are more frequently found at larger depths^23, 38^, the correlation coefficients of the mixed GABAergic populations would be expected to decrease with depth, as observed in our experiment. Finally, the ability of Bessel focus scanning to monitor the activity of inhibitory neuron ensembles in large volumes, makes it a powerful tool to elucidate how they shape the dynamics of cortical activity during learning^29^ as well as during distinct attentional and behavior states^24–27, 39–42^.

### 30 Hz volumetric imaging of spinal projection neurons in zebrafish larvae

In addition to mammalian brains, Bessel focus scanning can also be applied to other model systems. Even for optically transparent brains such as those of zebrafish larvae, where widefield imaging methods were used to image volumes at high rates^43, 44^, the Bessel approach still offers superior volumetric imaging performance, if the neural ensembles under investigation are distributed sparsely in 3D. One example is the set of spinal projection neurons in the hindbrain of zebrafish larvae. These neurons, which form a stereotyped array in which many cells are uniquely identifiable across individuals^45^, are responsible for the descending control of swimming behaviors, including sensory-evoked escapes^46^. While some of these neurons appear to be dedicated to controlling specific movements^46, 47^, there is evidence for modules that control different kinematic features across behaviors^48–50^, as well as for more distributed control of swim kinematics^51^. There does not yet exist an integrated understanding of how this population controls swimming. Since larval swimming occurs in discrete bouts separated by as little as 100 milliseconds, relating activity in this population to behavior will require imaging simultaneously across all the neuronal groups with high temporal resolution. We injected Cal520 dextran into the ventral spinal cord^52^ of *mitfa-/-* (nacre^53^) zebrafish. We then examined the calcium dynamics of the retrogradely labeled reticulospinal neurons in paralyzed larvae (7 days post fertilization) in response to a 4-ms mechanical and acoustic stimulus that induces fictive escape behavior (**Methods**). Unlike conventional imaging where neurons at different *z* planes had to be interrogated separately^51, 54^, a Bessel focus (NA 0.4, axial FWHM 64 μm) enabled us to image the reticulospinal neurons within a 270μm × 270 μm × 100 μm volume at 30 Hz (**Supplementary Video 4**). Due to minimal overlap between these neurons when viewed dorsally, all the neurons visible in the conventionally obtained *z* stack (**Fig. 4a**) were identifiable in the projected volume view offered by the Bessel focus scan (**Fig. 4b**). During nine repeated presentation of the stimulus (“tap” in **Fig. 4c**), most neurons showed stimulus-evoked calcium transients, consistent with previous recordings during escape behavior^51^. Additionally, bursts of activity were observed between stimulus presentations in sets of neurons that were distinct from, but overlapping with, those activated by the acoustic stimulus (**Fig. 4c**). The ability to monitor the activity of neurons located at different depths simultaneously makes it possible to detect trial-to-trial variations in the ensemble activity and discover neurons with correlated or anti-correlated activity patterns (**Fig. 4c**).

**Figure 4.**
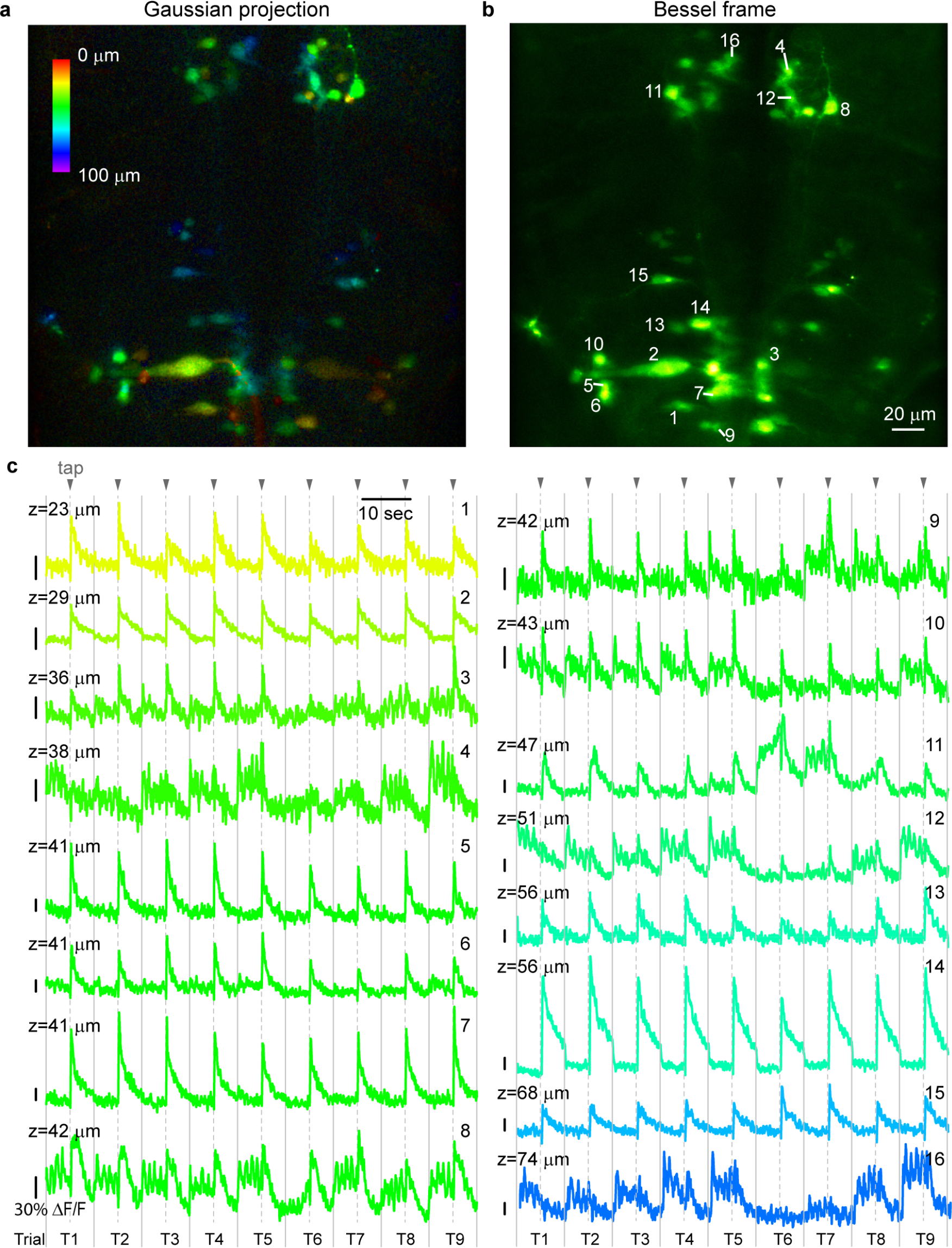
30 Hz volumetric imaging of spinal projection neurons in zebrafish larvae. (**a**) Mean intensity projection of a 3D volume (270μm×270μm×100μm, total 100 2D images) of retrogradely labeled reticulospinal neurons collected by scanning a Gaussian focus in the hindbrain of a zebrafish larva, with structures color-coded by depth. Post-objective power: 27 mW. (**b**) Image of the same neurons collected by scanning a Bessel beam with 0.4 NA and 64 μm axial FWHM. Post-objective power: 78 mW. (**c**) Calcium transients of individual neurons during 9 repeated presentation (“Trial” T1 to T9) of a 4-ms mechanical and acoustic stimulus (“tap“). Traces color coded by depth of neuron. Objective: Olympus 25×, 1.05 NA; Wavelength: 920 nm; Scale bar: 20 μm.

### Characterization of visually evoked responses in sparsely as well as densely labeled fruit fly brains

The fruit fly (*Drosophila melanogaster*) is an important model organism for the field of neuroscience^55^. The availability of sophisticated genetic tools and the small size of the fly brain have made it possible to record activity from anywhere inside the brain^56–58^. We first used Bessel focus scanning to image GCaMP6f^+15^ neurites in *UAS-GCaMP6f;33A12-GAL4*^59^ flies (see **Methods**). Presenting the same drifting grating stimuli described above to head-fixed flies (**Figs. 5a,b**), we monitored the calcium dynamics of a volume of neurites comprising most of the central complex (190 μm × 95 μm × 60 μm, **Fig. 5b**), a highly structured neuropil in the center of the fly brain, at a rate of 2.5 Hz, using Bessel focus (NA 0.4, axial FWHM 35 μm) scanning. The imaged volume contained fine neurites surrounding the ellipsoid body (a substructure of the central complex) (**Fig. 5c**), whose activity can be captured by a single Bessel frame (**Fig. 5d**). We found that the columnar neuron neurites within the ellipsoid body on either side of the central complex midline (ROIs 1 and 2 in **Fig. 5d**) had strongly anti-correlated activity (*R* = −0.73), which was not synchronized to the presentation of visual stimulation. In contrast, two neuronal processes above the ellipsoid body on both sides of the midline (ROIs 3 and 4) were strongly activated by drifting grating stimuli with no preference with respect to grating orientation (**Fig. 5f, Supplementary Video 5**).

**Figure 5.**
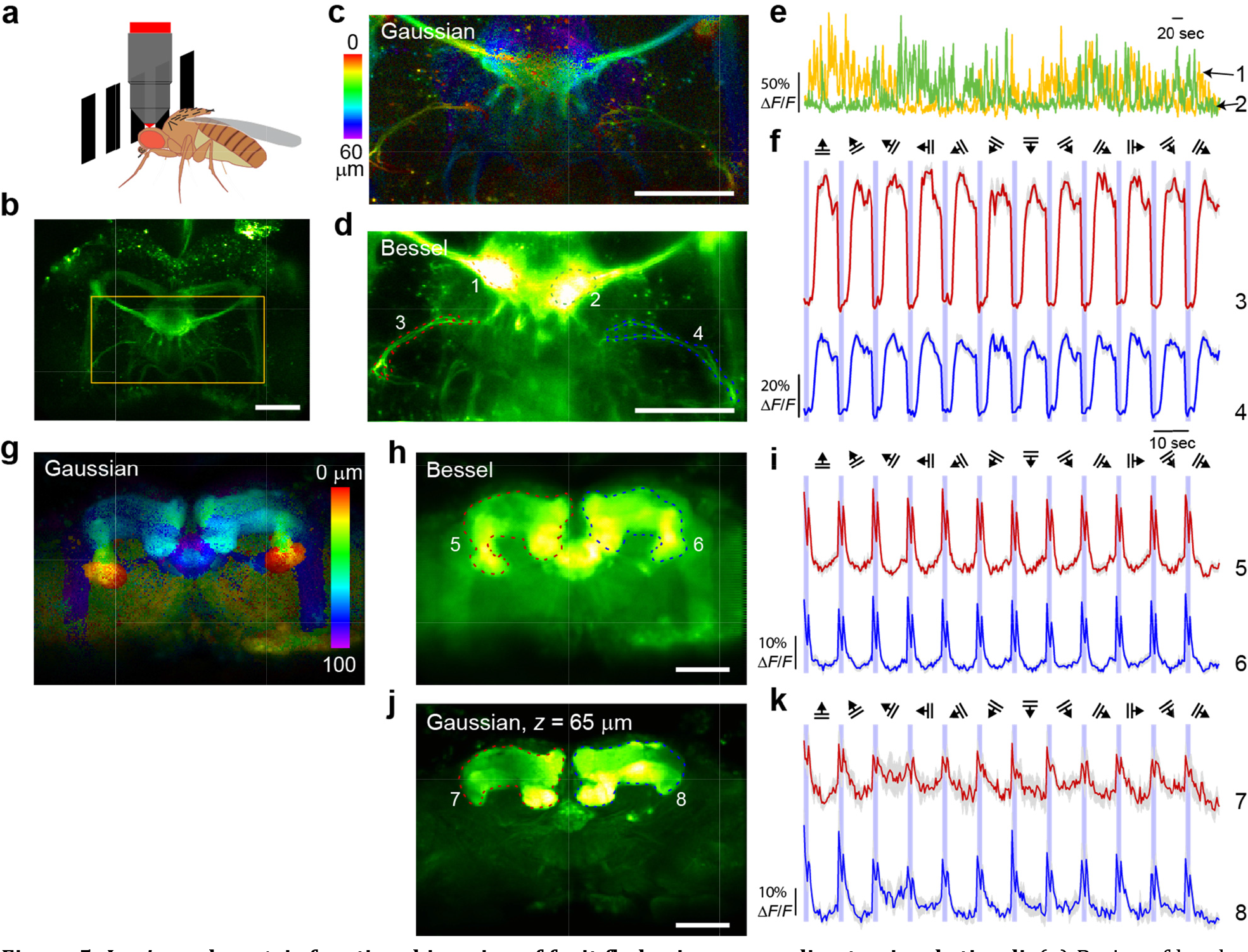
*In vivo* volumetric functional imaging of fruit fly brains responding to visual stimuli. (**a**) Brains of head-fixed flies were imaged during the presentation of grating stimuli. (**b**) *In vivo* image of the brain of a *UAS-GCaMP6f;33A12-GAL4* fly. Orange box indicates functional imaging volume. (**c**) Mean intensity projection of a 3D volume (190μm×95μm×60μm, total 61 2D images) of GCaMP6f+ neurites collected by scanning a Gaussian focus, with structures color-coded by depth. Post-objective power: 14 mW. (**d**) Image of the same neurites collected by scanning a Bessel beam with 0.4 NA and 35 μm axial FWHM. Post-objective power: 82 mW. (**e**) Neurites within the left and right lobes of the ellipsoid body (ROIs 1 and 2 in **d**) showed anti-correlated calcium activity. (**f**) 10-trial-averaged calcium transients of two neurites (ROIs 3 and 4 in **d**) strongly driven by grating visual stimuli. (**g**) Mean intensity projection of a 3D volume (307μm×216μm×100μm, total 101 2D images) of a *GCaMP6f;57c10-GAL4* fly with pan-neuronal expression of GCaMP6f, collected by scanning a Gaussian focus, with structures color-coded by depth. Post-objective power: 11 mW. (**h**) Image of the same brain collected by scanning a Bessel beam with 0.4 NA and 78 μm axial FWHM. Post-objective power: 31.5 mW. (**i**) Calcium transients (10-trial-averaged) from ROIs 5 and 6 in **h** were evoked by both the appearance of a static grating (data points within purple stripes) and the onset of its drifting movement (data points after purple stripes). (**j**) Optically sectioned image at z=65 collected by Gaussian focus scanning. Bright structures are the left and right lobes of the mushroom body. (**k**) Optically sectioned responses (10-trial-averaged) in mushroom body (ROIs 7 and 8 in **j**) shared similar temporal dynamics to the axially averaged responses in **i**. Objective: Olympus 25×, 1.05 NA; Wavelength: 940 nm; Scale bar: 50 μm.

We also applied Bessel focus scanning to fly brains with pan-neuronal expression of GCaMP6f *(UAS-GCaMP6f;57C10-GAL4*^59^). Scanning with an extended Bessel focus (NA 0.4, axial FWHM 78 μm) allowed us to measure visually evoked responses from a 307 μm × 216 μm × 100 μm volume (**Figs. 5g,h**), ~25% of the entire adult brain, at 3.6 Hz. Interestingly, the pixels with the strongest responses to grating stimuli in our imaging volume (**Figs. 5h,i, Supplementary Video 6**) fell within the area of the mushroom bodies, structures mostly known for olfactory learning and memory but not for visually driven responses in *Drosophila*. Because Bessel focus scanning cannot resolve structures along the *z* direction, we switched the microscope to conventional scanning with a Gaussian focus to confirm these responses. Placing the Gaussian focus within the mushroom bodies (**Fig. 5j**) and presenting the fly with the same grating stimuli, we again observed visually driven responses in the mushroom bodies (**Fig. 5k, Supplementary Video 7**). Both the axially averaged responses obtained via Bessel focus scanning and the optically sectioned responses obtained via conventional scanning shared similar features: calcium transients were evoked by both the appearance of new stationary gratings (data points within the purple stripes) and the drifting movement of the gratings (data points outside the purple stripes), and in both cases, the transients rapidly decayed with time. Our observation that mushroom bodies exhibit visually evoked responses is consistent with the recent discovery of Kenyon cells in the mushroom body that selectively responds to visual stimulation^60^. Given that the adult fly brain has a lateral extension (590 μm × 340 μm) that is within the typical field of view of 2PLSM and a thickness (120 μm) that is within the axial extend of our Bessel foci, Bessel focus scanning shows great promises as a method for rapid surveys of stimulus-or behavior-related neural activity in the entire fly brain.

## DISCUSSION

To understand information processing in neurons and neural networks that extend in three dimensions, we need to record neural activity with sub-second volume rate, while ideally maintaining synapse-resolving ability. Taking advantage of the fact that, during most *in vivo* brain imaging experiments, the structures of interest are stationary, we obtained volume information by scanning in 2D an axially extended focus approximating a Bessel beam. Using Bessel foci optimized to maintain high lateral resolution *in vivo*, we imaged the brains of a variety of species, including fruit fly, zebrafish, mouse, and ferret, and demonstrated high-speed volumetric imaging of neurons and neural compartments *in vivo*. The idea of extended depth-of-field imaging dates back 50 years^3^, and axially extended Bessel foci have been combined with 2PLSM to image brain slices^5, 6^. In our work, we systematically characterized the *in vivo* imaging performance of Bessel foci of different NAs and lengths, and discovered that low-NA Bessel foci provided optimal performance. Preserving sub-micron, synapse-resolving lateral resolution while reducing the background generated by the outer rings, these Bessel foci enabled us to measure calcium activity from single spines on multiple dendritic branches at up to 30 Hz, a combination of resolution and speed that is not matched by other volume imaging methods. Even though this performance came with the sacrifice of axial resolution, earlier studies^4, 6^ have shown that, given sufficient signal-to-noise ratio and a 2× reduction in volume imaging speed, axial resolution could be recovered during Bessel focus scanning through parallax-based stereo microscopy.

Now that we have discovered the optimal Bessel foci for *in vivo* extended depth-of-field imaging, a technical discussion on the best way to generate them is in order. The most common method to generate a Bessel focus is by using an axicon (conical lens). Compared to our module that employs a SLM, axicon-based systems are simpler^5, 6^, but can only generate a Bessel focus with fixed NA and axial length. For experiments requiring various combinations of NA and length, a SLM-based module is preferable, due to its flexibility and versatility. Furthermore, in addition to its ability to generate longer Bessel foci than axicons^4^, a SLM may also shape the axial intensity profile. For example, using a SLM, we generated a Bessel focus with higher intensity at larger depth, which improved image quality for deeper structures by compensating for the scattering loss of the excitation light (**Supplementary Fig. 7**).

Regardless of how a Bessel focus is generated, once incorporated into a microscope, the volume imaging speed depends on the lateral scanning speed of the microscope, with 30Hz volume rate at 512×512 lateral pixels easily achievable in systems equipped with resonant galvos or acousto-optic deflectors. With these systems, much higher volume speed can be achieved by simply reducing the number of lateral pixels. Ultimately, with heating and photodamage of the brain placing an upper limit on excitation power^61^, the practically achievable volume imaging speed depends on the fluorescence probe, with brighter probes requiring less dwell time at each pixel and thus enabling higher imaging speed.

What is the limit on the axial extent of the volume? The longest Bessel focus that we generated to image brains *in vivo* was 400 μm at 0.4 NA for the Olympus 25× objective (data not shown). Even though longer Bessel foci may be generated by the SLM, the practically useful axial length is determined by both the sample and the desired lateral resolution. Sparsely labeled samples have lower probability of having structures overlap in their *xy* locations, thus allow longer foci to be used. However, at the same NA, longer Bessel foci have more energy in the side rings surrounding the central focus, which may lead to higher background signal (**Supplementary Fig. 1**). Relaxing the lateral resolution requirement (e.g., from submicron, synaptic resolution to cellular, micron-level resolution) would allow lower-NA Bessel foci to be employed, where background contamination by the side rings is less of an issue. Another practical limitation on the length of the Bessel focus is optical power. In general, compared to Gaussian focus scanning, the redistribution of power into the side rings during the formation of a Bessel focus requires the average excitation power to be increased. Care needs to be taken to ensure that the physiology of the brain is not altered by the deposited laser power^61^.

**Supplemental Figure 7.**
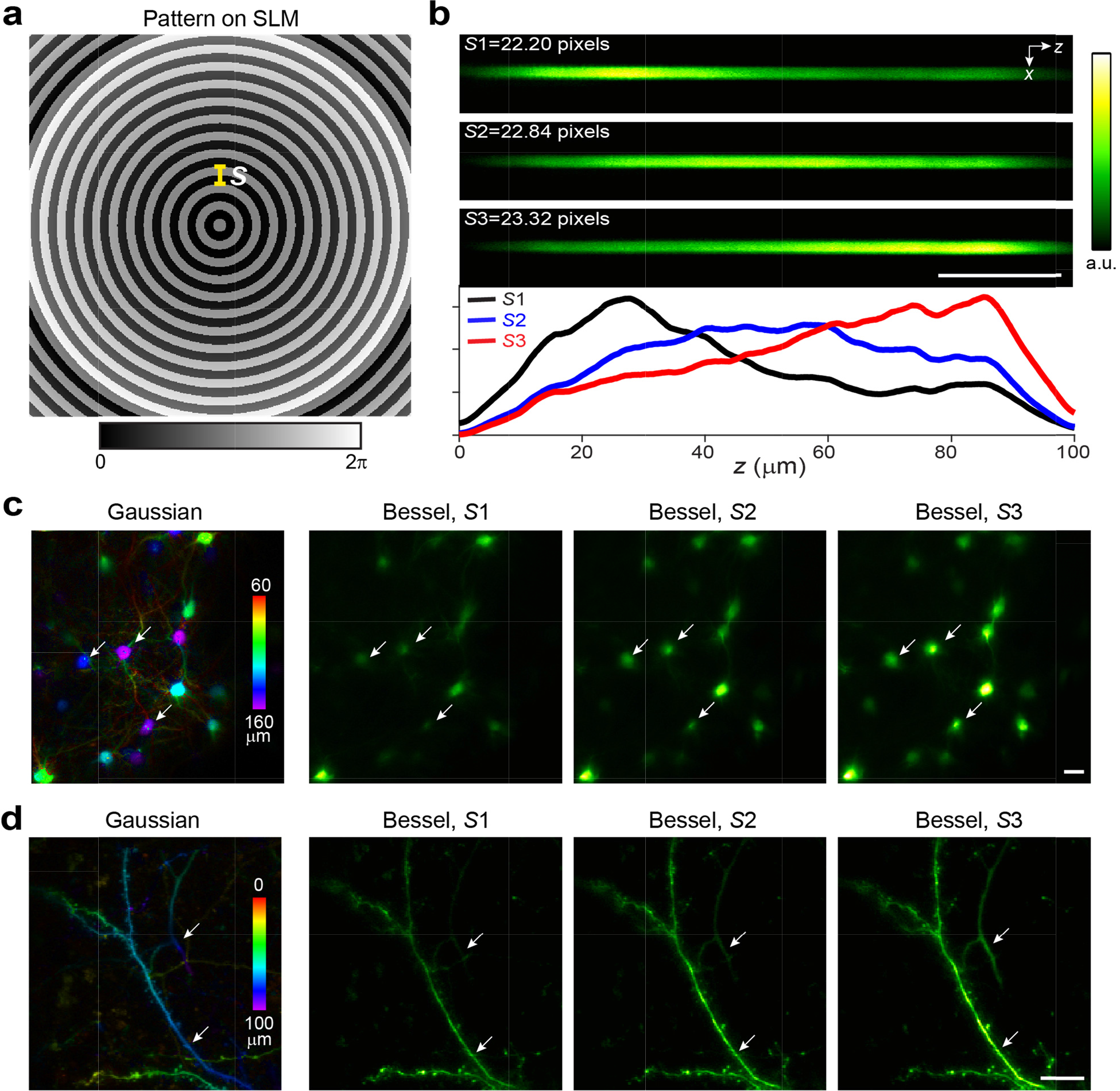
Engineering Bessel foci with asymmetric axial intensity distribution with a SLM. (**a**) Overlaying a binary concentric ring pattern on a quadratic phase pattern on the SLM allows the shaping of axial intensity distribution of the Bessel focus. *S*: period of the concentric rings in units of pixels. (**b**) Axial images and corresponding intensity profiles of a 2-μm-diameter bead using three different Bessel foci generated with periods *S*1, *S*2, *S*3. (**c,d**) (From left to right) *in vivo* images of neurons and neurites measured in V1 of awake mice using Gaussian focus, Bessel foci generated with *S*1, *S*2, and *S*3, respectively. Gaussian images are mean intensity projections with features color-coded by depth. Arrows point to deeper structures that had best image quality when the Bessel focus of increasing intensity with depth (*S*3 in **b**) was used. Objective: Olympus 25×, 1.05 NA; Post-objective power: **c**, 14 mW for Gaussian modality and 32 mW for Bessel modality; **d**, 21 mW for Gaussian modality and 61 mW for Bessel modality; Wavelength: 940 nm; Scale bars: 20 μm.

The application of Bessel focus scanning is not limited to neurons. Any sample with sparsely labeled stationary structures can be imaged at high volume rates using this approach. For densely labeled brains, Bessel scanning can be used as a high-throughput screening method to identify areas of activity followed by investigations at higher axial resolution (e.g., **Figs. 5g-k**). For studies of densely labeled brains where single neuron resolution is required, Bessel scanning may also be applied, if the activity pattern is sparse and spatially overlapping calcium responses can be unmixed temporally^62, 63^.

In addition to the fact that video-rate volume imaging with synaptic resolution, as reported here, has not been demonstrated by other methods, Bessel focus scanning has several additional advantages when compared to volume imaging methods that rapidly move a Gaussian focus in 3D^64^. By projecting 3D volume information into 2D, Bessel focus scanning leads to ten to hundred fold reduction in data size. Together with its intrinsic insensitivity to sample *z* motion, this extended depth-of-field approach to volume imaging substantially reduces the amount of data and complexity of data analysis. In addition, Bessel focus scanning can be integrated into existing microscope systems without any change to the data acquisition software. (In our experiments, Bessel modules were added to three systems run by three different control software – homemade, open source^65^, and commercial, respectively). Finally, the Bessel module can be combined with existing scanning hardware (e.g., galvos, acousto-optical deflectors, fast axial scanning units) and modes (e.g., raster scanning, line scanning, random accessing)^64^ to achieve higher speed, larger volume, and more versatile experiments to study brains in action.

## METHODS

All animal experiments were conducted according to the National Institutes of Health guidelines for animal research. Procedures and protocols on mice and zebrafish were approved by the Institutional Animal Care and Use Committee at Janelia Research Campus, Howard Hughes Medical Institute. Experimental protocols on ferret were approved by the Max Planck Florida Institute for Neuroscience Institutional Animal Care and Use Committee.

### Assembly, incorporation, and control of Bessel modules in three 2PLSMs

A focus approximating a Bessel beam can be generated by illuminating the back pupil plane of a microscope objective with an annular pattern (i.e., a ring). We chose to use a reflective phase-only spatial light modulator (SLM, CODPDM512-800-1000-P8, Meadowlark Optics) to generate annular illumination, because it allowed us to produce Bessel foci of different NAs and axial lengths and systematically test their imaging performance *in vivo* (**Supplementary Fig. 1**). With the SLM placed at the front focal plane of lens L1, an electric field distribution that was the Fourier transform of the phase pattern on the SLM was generated at the back focal plane of L1^66^, where we placed an annular apodization mask. The apodization mask was then imaged onto the galvos and the back pupil plane of the microscope objective using pairs of lenses (L2 and L3, L4 and L5) in 4*f* configuration (**Fig. 1b**). For ease of alignment, the SLM was mounted on a platform with 3D translational and two-axis rotational controls, and the annular mask was mounted on a 3D translational stage. When incorporating Bessel modules into two homebuilt 2PLSM in Janelia and one commercial 2PLSM (Thorlabs) in Max Planck Florida, different lenses were used to match the magnification factor and objective used in each system (**Supplementary Table 1**).

A standalone control program (Meadowlark Optics) displayed phase patterns on the SLM, with no communication with or modification of the microscope software needed. Concentric binary grating patterns on the SLM were used to generate annular illumination at the apodization mask^4^ (see **Supplementary Notes** and **Supplementary Codes** for details). By selecting a specific annulus on an apodization mask array (Photo Sciences, Inc.) to truncate the electric field distribution at the back focal plane of L1, we can fine-tune the axial lengths and intensity distributions of the Bessel-like foci (**Supplementary Fig. 8**). (Strictly speaking, a Bessel beam is generated by an infinitely thin annulus, has infinite axial length, and carry infinite energy. The Bessel-like foci generated here with annuli of finite widths are known as quasi-Bessel, or finite Bessel beams.) In general, the NA of a Bessel focus is dictated by the outer diameter of the annulus, while its axial length additionally depends on the thickness of the annulus, with thinner annuli generating longer Bessel foci (but higher background). The combination of SLM with the apodization mask array enabled independent adjustments of the NA and axial length of the Bessel focus. Overlaying a defocus pattern on the binary concentric grating, we engineered Bessel foci with higher axial intensity at larger *z* depth to facilitate the imaging of deeper structures in scattering brains (**Supplementary Fig. 7**).

**Supplementary Figure 8.**
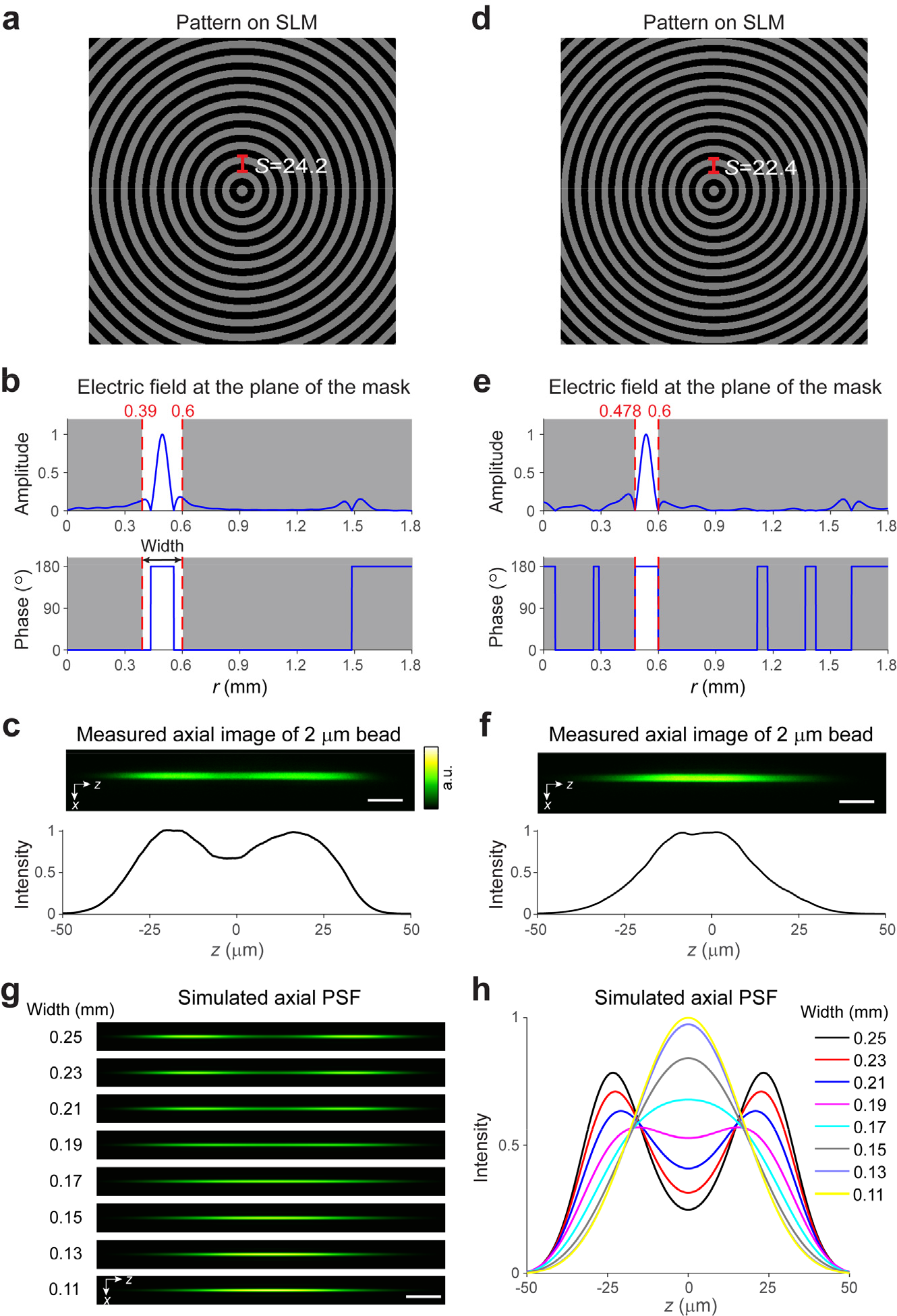
Annular apodization mask fine-tunes the axial length and intensity distribution of Bessel foci. (**a-c**) and (**d-f**): two NA-0.4 Bessel foci with different axial lengths and intensity distributions. (**a,d)** Binary concentric grating patterns (phase values: 0 and π) on SLM. *S* indicates the period of the gratings in units of pixels. (**b,e**) (Top) amplitude and (bottom) phase of the electric field of the light at the plane of the annular apodization mask (0.39 mm inner radius and 0.6 mm outer radius for **b**, 0.478 mm inner radius and 0.6 mm outer radius for **e**). Electric field within the annulus (areas between red dashed lines) was transmitted. (**c,f**) (Top) axial (*xz*) images of 2-μm-diameter fluorescent beads and (bottom) line intensity profiles along the *z* axis obtained with Bessel foci generated by **a** and **b**, **d** and **e**, respectively. (**g**) Simulated axial PSFs and (**h**) their intensity profiles along *z* as a function of annulus width of the apodization mask. Incidence beam diameter at SLM: 2.2 mm. Magnification from mask to objective back pupil: 4.7; Objective: Olympus 25×, 1.05 NA; Post-objective power: 14.2 mW and 15.5 mW for **c** and **f**, respectively; Wavelength: 900 nm; Scale bar: 10 μm.

### Mouse preparation

Mice (females or males, >2-months-old) were housed in cages under reverse light cycle. Wild-type mice were used for all mouse experiments except those in **Fig. 3**, where Gad2-ires-Cre (Jackson Laboratories, Gad2tm2(cre)Zjh/J, stock #: 010802), SOM-ires-Cre (Jackson Laboratories, Sst<tm2.1(cre)Zjh>/J, stock # 013044), and VIP-ires-Cre (Jackson Laboratories, Vip<tm1(cre)Zjh>/J, stock # 010908) mice^23^ with Cre-dependent GCaMP6s virus injection (AAV2/1.syn.Flex.GCaMP6s, ~8 × 10^11^ infectious units per ml) in V1 were used. Sparse expression of GCaMP6s was achieved by injecting a mixture of diluted AAV2/1-syn-Cre particles (original titer: ~10^12^ infectious units per ml, diluted 8,000-fold in phosphate-buffered saline) and high titer, Cre-dependent GCaMP6s virus (AAV2/1.syn.Flex.GCaMP6s, ~8 × 10^11^ infectious units per ml).

Virus injection and cranial window implantation procedures have been described previously^67^. Briefly, a 2.5-mm diameter craniotomy was made over either V1 or S1 of mice that were anaesthetized with isoflurane (1-2% by volume in O_2_) and given the analgesic buprenorphine (SC, 0.3 mg per kg of body weight). Dura was left intact. Virus injection was then performed using a glass pipette (Drummond Scientific Company) beveled at 45° with a 15-20-μm opening and back-filled with mineral oil. A fitted plunger controlled by a hydraulic manipulator (Narishige, M010) was inserted into the pipette and used to load and slowly inject 10-20 nl viral solution into the brain (~200-400 μm below pia). After the pipette was pulled out of the brain, a glass window made of a single coverslip (Fisher Scientific no. 1.5) was embedded in the craniotomy and sealed in place with dental acrylic. A titanium head-post was then attached to the skull with cyanoacrylate glue and dental acrylic. *In vivo* imaging was carried out after two weeks of recovery and habituation for head fixation. All imaging experiments were carried out on headfixed awake mice, except data in **Fig. 2**, where mice were anesthetized to reduce calcium transients induced by BAP.

### Ferret preparation

Juvenile female ferrets (*Mustela putorius furo*, Marshall Farms) housed in a vivarium under 16h light/ 8h dark cycle were used. Sterile surgical procedures have been described previously^16^. Briefly, virus injection was carried out in P21-P23 juvenile female ferrets anesthetized with ketamine (50 mg/kg, IM) and isoflurane (1-3% in O_2_), then intubated and artificially respirated. Atropine (0.2 mg/kg, SC) was administered to reduce secretions. 1:1 mixture of lidocaine and bupivacaine was administered subcutaneously in the scalp. With the internal temperature maintained at 37°C with a heating pad, a 0.8-mm-diameter craniotomy was made over ferret V1. AAV2/1.hSyn.Cre (1.32×10^13^ infectious units per ml, Penn Vector Core) was diluted by 100,000 fold in PBS (Sigma) and mixed with AAV2/1.CAG.Flex.GCaMP6s (7.31×10^12^ infectious units per ml, Penn Vector Core) to sparsely express GCaMP6s in layer 2/3 cortical neurons. Beveled glass micropipettes (~15 μm outer diameter) were lowered into the brain, and 55 nl of virus were injected (Nanoject II, Drummond Scientific Company) at 250 and 400 μm below the pia. To prevent dural regrowth and attachment to the arachnoid membrane, the craniotomy was filled with 1% w/v agarose (Type IIIa, Sigma-Aldrich).

After two to three weeks of expression, ferrets were anesthetized with ketamine and isoflurane and administered Atropine and bupivacaine, placed on a heating pad, then intubated after tracheotomy and artificially respirated. An intravenous cannula was placed to deliver fluids. ECG, endtidal CO_2_, external temperature, and internal temperature were continuously monitored during the procedure and subsequent imaging session. After scalp retraction, a titanium headplate was adhered to the skull using C&B Metabond (Parkell). A 5.5-to-6.0-mm diameter craniotomy was performed at the viral injection site and the dura retracted to reveal the cortex. A 5.0-mm outer diameter titanium cannula with a glass bottom (affixed with optical adhesive # 71, Norland Products) was placed onto the brain to gently compress the underlying cortex and dampen brain motion. The cranial cannula was hermetically sealed using a stainless steel retaining ring (5/16” internal retaining ring, McMaster-Carr), Kwik-Cast (World Precision Instruments), and Vetbond (3M). A 1:1 mixture of 1% Tropicamide Ophthalmic Solution (Akorn) and 10% Phenylephrine Hydrochloride Ophthalmic Solution (Akorn) was applied to both eyes to dilate the pupils and retract the nictating membranes. Eyes were lubricated hourly with Silicon Oil AP 150 Wacker (Sigma-Aldrich). Upon completion of the surgical procedure, Isoflurane was gradually reduced and then vecuronium (2 mg/kg/hr) or pancuronium (2 mg/kg/hr) was delivered Intravenously to immobilize the animal.

### Zebrafish preparation

A 25% w/v solution of Cal520-Dextran conjugate (3000MW, AAT Bioquest, California) in zebrafish external solution (134 mM NaCl, 2.9 mM KCl, 2.1 mM CaCl_2_, 1.2 mM MgCl_2_ and 10 mM HEPES glucose, pH 7.8^68^) was pressure injected into the ventral spinal cord of 6 day post fertilization *mitfa-/-* (nacre^53^) fish larvae just anterior to the anal pore, following standard procedures^52^. The larvae were left in E3 embryo medium (5mM NaCl, 0.17mM KCl, 0.33mM CaCl_2_ and 0.33mM MgSO_4_) at 28°C overnight to allow retrograde filling of reticulospinal neurons. On the next day before imaging, larvae were paralyzed by bath application of 1 mg/ml of alpha-bungarotoxin (Molecular Probes) in embryo medium for 1 min. Larvae were mounted dorsal-side up on dishes coated with Sylgard 184 (Dow Corning) in a small drop of 1.5% low melting point agarose (Invitrogen), covered with embryo medium. During imaging, the dish containing the larval zebrafish was mounted on top of an acrylic platform with an attached speaker that delivered acoustic and mechanical stimuli. The stimulus, which consisted of a 500 Hz sinusoidal waveform lasting 4 ms, was delivered every 100 seconds. Images were acquired for 5 s before and after the stimulus, which was repeated 9 times.

### Fly preparation

Flies were prepared for two-photon imaging as previously described^56^. Briefly, flies were anesthetized on ice and transferred to a cold plate at 4°C. They were glued to a stick with UV glue and inserted into an imaging holder (made from a stainless steel shim using a laser mill) with micromanipulators under a dissection stereomicroscope. The fly’s head and a part of the thorax were glued to the holder using UV gel. The holder isolated the back of the fly’s head and a small part of the thorax for imaging while leaving the fly’s eyes fully exposed for visual stimulation. The cuticle on the backside of the fly’s head was removed with dissection needles and forceps and the brain of the fly was exposed for two-photon imaging. To minimize brain movement, we immobilized the fly’s proboscis by pressing it into the fly’s head and fixing it with wax. We additionally cut muscle M16 or the nerves innervating the muscle. The holder with the fly was then transferred to the microscope for two-photon imaging.

### Visual stimulation for the mouse and the fly

UV (320-380 nm, peak at 365 nm, Richard Wolf) and visible (450–495 nm, peak at 472 nm, SugarCUBE) light was used to evoke activity in the fly and the mouse brains, respectively. Visual stimuli were presented by back projection on a screen made of Teflon^®^ film (McMaster-Carr) using a custom-modified DLP projector^69^. For mouse experiments, the screen was positioned 17 cm from the right eye, covering 75° × 75° degrees of visual space and oriented at ~40° to the long axis of the animal. Full-field square gratings were presented for 12 seconds (static for the first and last 2.5 sec and moving during the middle 7 sec) at 12 orientations (0° to 330° at 30° increments) in pseudorandom sequences. A total of ten trials were repeated in each measurement. For some experiments, blank screens with equal mean luminance were presented between grating stimuli. Only data collected during grating movements were used to calculate orientation tuning curves. Gratings had 100% contrast, 0.07 cycles per degree, and drifted at 26 degrees per sec (2 Hz temporal frequency). The same stimulus protocol was used for the fly, except that the body axis of the fly was orthogonal to the screen. During mapping of single-spine calcium transients, gratings with reduced contrast, together with anesthesia, were used to minimize the firing of somatic action potentials^15^.

### Image processing and analysis

Imaging data were processed with custom programs written in MATLAB (Mathworks) or Fiji^11^. Images were registered either with an iterative cross-correlation-based registration algorithm^67^, or with TurboReg plugin in Fiji. ROIs were selected and outlined manually. The averaged fluorescent signal within the ROI was used to calculate calcium transients. For GCaMP6s^+^ inhibitory neuron experiments (**Fig. 3**), neuropil background was subtracted from averaged signal before calculating calcium transients (**Supplementary Fig. 5**).

For experiments that characterized the tuning of dendritic spines (**Fig. 2**), BAP-induced calcium signal in the spine was removed from the measured transients as described previously^15, 16^. Briefly, BAP-induced signal was calculated from a ROI covering the entire dendritic shaft in the image. This global dendritic signal was then removed from the spine signal using a robust regression method (robustfit function in MATLAB). The response *R* of each spine to a visual stimulus was defined as the average Δ*F/F* across the 7-s window of drifting gratings. The distribution of *R* was assumed to be normal and variances were assumed to be equal across grating angle *θ* but this was not formally tested. A spine was defined as orientation selective if its responses across the various drifting directions were significantly different (ten trials, one-way ANOVA, P < 0.01). For these spines, we determined their preferred grating drifting angle *θ*_pref_ by fitting their normalized response tuning curves (e.g., **Fig. 2d, Supplementary Fig. 4c**) with a bimodal Gaussian function^70^:

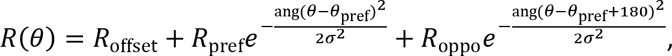

where *R*_offset_ is a constant offset, and *R*_pref_ and *R*_oppo_ are the responses at *θ*_pref_ and *θ*_pref_-180 degree, respectively. The function ang(*x*) = min(|x|, |x − 360|, |x + 360|) wraps angular values onto the interval 0° to 180°.

## ACKNOWLEDGEMENTS

The authors thank Vivek Jayaraman for helpful discussion and fly lines, Yoshi Aso and Yi Sun for help with fly anatomy, and Julie Kuhl for help with illustration. R.L., W. S., Y. L., A. K., J. S., B. M., M. T., M. K., and N. J. were supported by Howard Hughes Medical Institute. M. T. was also supported by Japanese Society for the Promotion of Science (S2602 and 15K06708). D.E.W., B.S., and D.F. were supported by National Eye Institute of the NIH under Grant 2 R01 EY011488-17. J.B. was supported by a PhD fellowship from the Portuguese Fundação para a Ciência e a Tecnologia. M.B.O was supported by the Champalimaud Foundation and the Bial Foundation (185/12).

### CONTRIBUTIONS

N.J. conceived and oversaw the project; R. L. and N.J. designed the Bessel modules; R.L. and N.J. built Bessel modules with help from A.K., W.S., B.M., D.W., and B.S.; W.S. and Y.L. prepared mouse samples; R.L., W.S., Y.L., A.K., B.M. and N.J. collected mouse data; M.K. and M.B.O. designed zebrafish experiments; J.B. prepared zebrafish samples with help from M.T.; R.L., A.K., J.B., and N. J. collected zebrafish data; J.S., R.L., N.J. designed drosophila experiments; J.S. prepared drosophila samples; R.L., J.S., and N.J. collected drosophila data; D.F. and N.J. designed ferret experiments; D. E. W. and B.S. prepared ferret samples and collected ferret data. All authors contributed to data analysis and presentation. R.L. and N.J. wrote the paper with inputs from all authors.

